# The Interplay Between Nutrition and the Dynamics of the Midgut Microbiome of the Mosquito *Aedes aegypti* Reveals Putative Symbionts

**DOI:** 10.1101/2024.03.04.583003

**Authors:** João Felipe M. Salgado, Balakrishnan N. V. Premkrishnan, Elaine L. Oliveira, Vineeth Kodengil Vettath, Feng Guang Goh, Xinjun Hou, Daniela I. Drautz-Moses, Yu Cai, Stephan C. Schuster, Ana Carolina M. Junqueira

## Abstract

Blood meals are crucial for the reproductive cycle of *Aedes aegypti* and represent the means by which arboviruses are transmitted to its hematophagy hosts. It also has been postulated that feeding on blood may modulate the mosquito microbiome, but the compositional shifts in microbial diversity and function remain elusive. In this paper, we analyzed the modulation of the midgut microbiome in 60 females of *Aedes aegypti* throughout the digestive period, 12, 24, and 48 hours after blood or sugar meals using whole-genome shotgun sequencing. Microbial transstadial transmission between larvae and adults was also assessed. This approach provided a high coverage of the midgut metagenome, allowing microbial taxonomic assignments at the species level and gene-based functional profiling. Females at later hours post-feeding and larvae display low microbiome diversities and little evidence of transstadial transmission. However, a striking proliferation of Enterobacterales was observed during early hours of digestion in blood-fed mosquitoes. The compositional shift was concomitant with a predicted functional change in genes associated with carbohydrate and protein metabolism. The observed shifts in blood-fed females’ midguts are restored to a sugar-fed-like microbial profile after 48h, when blood digestion is completed. Conversely, as in all blood-fed females, a high abundance of the opportunistic human pathogen *Elizabethkingia anophelis* (Flavobacteriales) takes place in this post-digestion stage. This bacterial species has also been described as a symbiont of mosquitoes of the genus *Anopheles* (Culicidae). This work is the first report of the adaptation of the midgut microbiome *of A. aegypti* to a digestive role after a blood meal, at the expense of the proliferation of potential symbionts.

**Significance statement:** The findings in this paper can contribute to a better understanding of the dynamics of the mosquito microbiome during digestion and its potential implications for host physiology and metabolism, also informing the future development of sustainable methods for insect-borne diseases control based on microbial components that might influence vectorial capacity and pathogen transmission by *A. aegypti*.

## 1. Introduction

The knowledge accumulated in the last decade about the association of multicellular eukaryotes with microorganisms has provided a paradigm shift in what was previously known of metabolic, physiological, and homeostatic fitness in virtually every multicellular host organism (Heiss and Olofsson, 2019; Kang et al., 2019). Microbiome research has primarily focused on humans, but recent advances expanded the investigation to other animals. In this context, several insect vectors have been targeted by metagenomic approaches, unveiling complex microbiome interactions that regulate processes essential to the host life cycle (Angleró-Rodríguez et al., 2017; Cappelli et al., 2019; Junqueira et al., 2017; Rodríguez-Ruano et al., 2018; Vivero et al., 2019). While diverse environmental factors may contribute to variations in the microbiome of insects (Junqueira et al., 2017; Saab et al., 2020), the nutritional source appears to be especially impactful in modulating microbial communities (Mason and Raffa, 2014; Yun et al., 2014). Likewise, the patterns and dynamics of microbial diversity in holometabolous insects can also be determined by their habitats and developmental stages (Douglas, 2015).

In hematophagous mosquitoes who rely on the blood meal for egg development, oviposition, and lifespan (Gaio et al., 2011; Petersen et al., 2018), the food type and source proved to be fundamental for the dynamics of their intestinal microbiome, influencing the vector’s susceptibility to viruses and their capacity to transmit pathogens to hosts (Almire et al., 2021; Apte-Deshpande et al., 2012; Ramirez et al., 2014; Sharma et al., 2013). These findings are essential for developing successful strategies for vector control based on microbiota manipulation, such as those reported for Wolbachia infections (Wasi et al., 2019; Aliota et al., 2016). The yellow fever mosquito, *Aedes aegypti*, is the primary vector of viral diseases worldwide, such as yellow fever, Zika, chikungunya, and dengue. The latter, alone, is responsible for a global economic burden of US$ 9 billion per year (Shepard et al., 2016; Shragai et al., 2017). Previous studies based on 16S sequencing have reported a core microbiome composed of aerobic and facultative-anaerobic bacteria in *Aedes* spp. (Scolari et al., 2019). However, multiple factors modulate *A. aegypti* microbiota, such as habitat, environmental contamination with fertilizers or antibiotics, sex, developmental stage, or nutrition (Scolari et al., 2019). Significant differences in the bacterial composition and diversity were found in the midgut of *A. aegypti* fed on distinct food sources and in mosquitoes fed with blood from different animal hosts (Muturi et al., 2021, 2018). Despite the critical role of hematophagy for *A. aegypti* reproduction, the microbial shifts triggered by the blood meal in the midgut and its modulation throughout the digestion have never been tackled by large-scale metagenomic approaches (Hyde et al., 2020).

In the present study, we provide for the first time an in-depth investigation of 70 individual metagenomes of *A. aegypti* performed by whole-genome shotgun sequencing of blood and sugar-fed mosquitoes. The microbiome dynamics were followed in both diet types in adults at 12h, 24h, and 48h after feeding. Our metagenomic approach allowed for the microbial taxonomic assignment up to the species level, revealing a large amount of Enterobacteria in the mosquito’s midgut during the blood meal digestion. This compositional shift was accompanied by a highly correlated functional change in microbial taxa involved in the catabolism of amino acids, sugar, and virulence in mosquitoes fed with blood. The sugar-fed group presents a significantly higher diversity in its microbiome when compared to the blood-fed group, but no significant functional correlations. The post-digestion period is associated with the increase of the bacterial species *Elizabethkingia anophelis* in both groups. This study is the first report of the occurrence of this symbiotic Flavobacterium in *A. aegypti* and its modulation in the gut microbiome caused by blood meals. Transstadial-transmitted microorganisms were also evaluated by comparing larval and adult microbiomes, showing no evidence for such phenomenon. Larvae displayed a low microbial diversity, with the predominance of the genus *Microbacterium*.

## 2. Results and discussion

### 2.1. Metagenomic datasets

The total DNA extraction of the midgut of 60 females of *A. aegypti* provided an average of 1.078 ng/µL ± 0.663, while ten individual larvae in the fourth instar (group L4) yielded 0.599 ng/μL ± 0.121. Adult mosquitoes fed with blood (group AB) showed a lower DNA yield when compared to the mosquitoes fed with sugar (group AS), with an average of 0.783 ng/μL ± 0.385 for AB and 1.372 ng/μL ± 0.752 for AS. The total DNA amount recovered from each individual sample is shown in Supplementary Figure 1 and Table S1. Despite the low biomass, the average number of reads generated per sample was approximately 45 million (negative and environmental controls excluded, Table S1). A total of 3,213,701,600 reads were generated, from which 1,376,183,852 reads (∼42.82%) were classified as non-host reads (NH) after *in-silico* removal of mosquito genomic sequences. The total DNA and metagenomic reads per group before and after mapping and the reads generated for controls are in Supplementary Table S1. A total of 1,164,457,880 paired-end reads (an average of: L4 = 38,054,610 ± 6,105,856; AB = 20,139,425 ± 11,488,999; AS = 5,990,967 ± 624,586) were used in the subsequent metagenomic analyses.

To assess whether the number of microbial phylotypes in each sample was a function of the sequencing depth, we performed interpolation and extrapolation of the reads using microbial richness at the genus and species taxonomic levels. The groups L4 and AB reach a *plateau* of rarefaction at approximately 400,000 reads. In comparison, samples from the AS group reached the rarefaction threshold at approximately 50,000 reads, showing that all curves were rarefied to the point where the size of the datasets no longer contributed significantly to the increase of microbial diversity (Supplementary Figure S2). Therefore, the metagenomic dataset sizes used in this work were not creating a bias in the diversity of the microbiomes.

### 2.2. Larval microbiome and experimental controls

Larvae are exclusively colonized by Actinobacteria of the genus *Microbacterium* (3,403,719 reads; 65 phylotypes detected; Figure 1). They display a significantly higher diversity with the Chao1 index (Figure 2A) when compared to the adult groups, but this result is due to a decreased evenness, which is reflected in the diversity analysis with Simpson reciprocal indices (Figure 2A). Recently, the role of *Microbacterium* sp. was assessed in axenic *A. aegypti* larvae, showing that it is the only taxon that was not associated with an increased rate of survival to adulthood (Coon et al., 2014). This finding suggests that *Microbacterium* sp. do not contribute significantly to the development of larvae, despite their dominance. Nevertheless, its colonization in *A. aegypti* larvae may be relevant in other aspects, including symbioses that antagonize the establishment of fungal entomopathogens, such as *Metarhizium robertsii* (Noskov et al., 2021). This perspective may explain why fungal pathogens such as *M. majus* and *Aspergillus flavus* coincide with low *Microbacterium* sp. colonization in adult mosquitoes. Additionally, the water sample where larvae were reared did not show the presence of *Microbacterium* sp. (Supplementary Figure S4), further indicating that its dominance in larval samples is not acquired from the environment and is indeed typical of this developmental stage. The Actinobacteria *Leifsonia aquatica* (111,018 reads)*, Leucobacter chironomi* (160,140 reads)*, and Corynebacterium sp.* (223,942 reads) are also found in larvae but not in adult mosquitoes. These results corroborate the hypothesis that there is little-to-no transstadial transmission of microbiome components, with considerable differences in compositional and diversity patterns from larval to adult mosquitoes.

**Figure 1.**
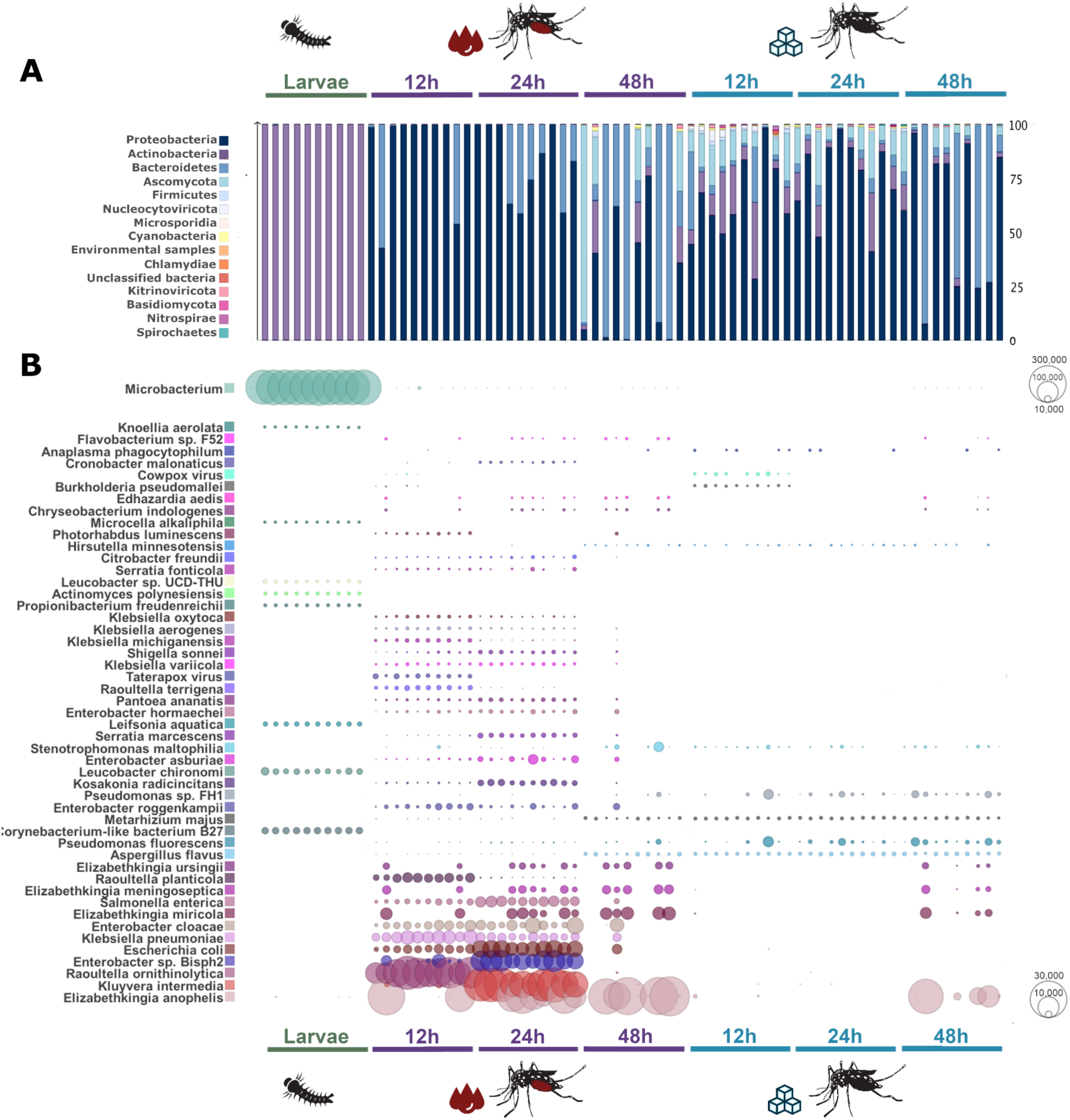
The microbiome composition of female adults and larval stages at phylum (A) and species (B) levels. Bubbles display the relative abundance of microbial species in quadratic scale based on the normalized number of assigned reads for each adult midgut or larval sample. The bins in the bar chart are scaled percentually. *Microbacterium* sp. were collapsed to the taxonomic level of the genus and displayed in its own scale (top-left) to enable the comparison between different taxa in the experimental groups.

**Figure 2.**
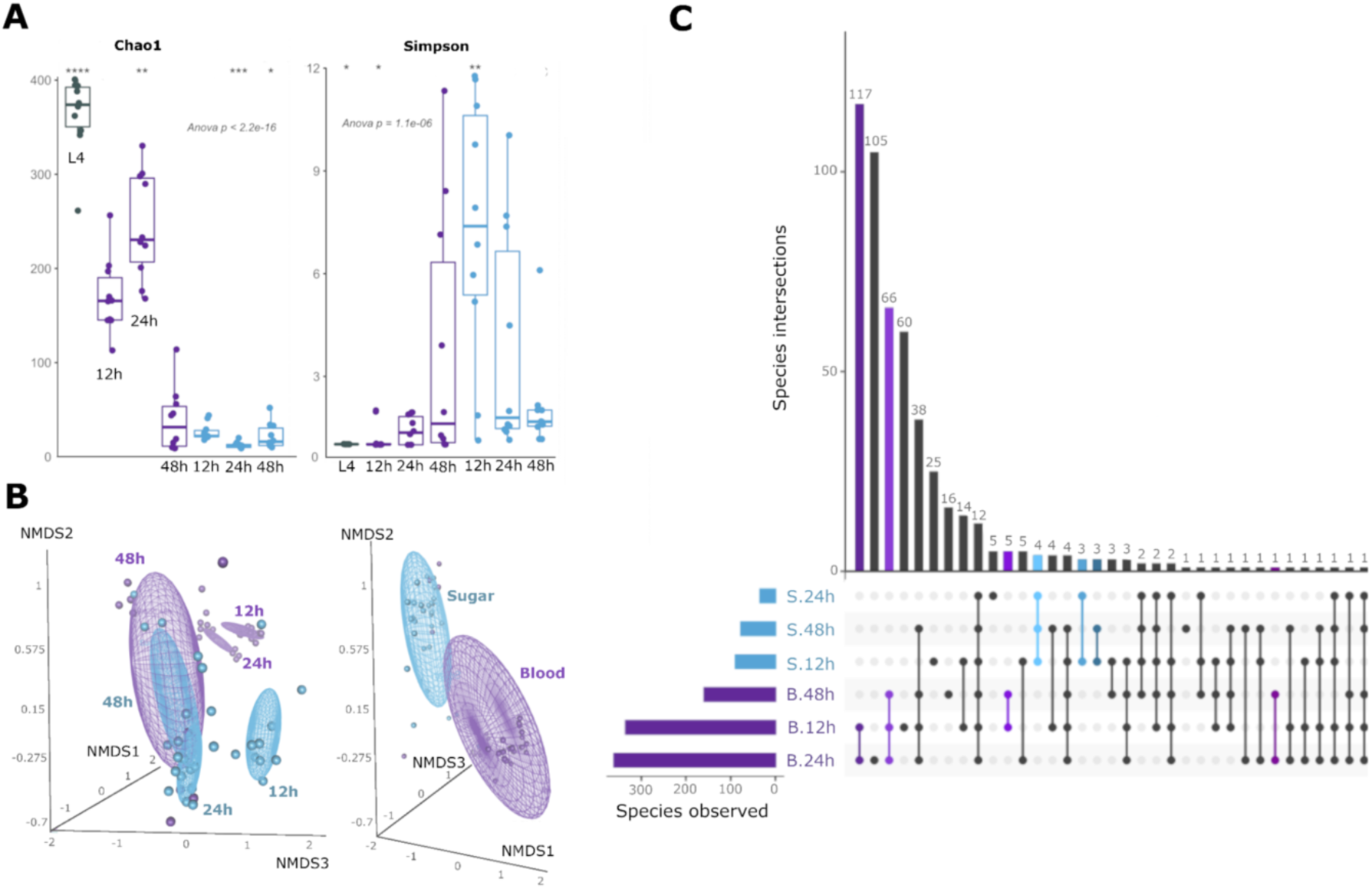
Diversity profiles of mosquitoes’ microbiome in different developmental stages and throughout 48 h of the digestion of different diets in adults. A) Boxplots showing the distribution of different diversity indices and entropies. The global and pairwise significances are assessed with ANOVA and Wilcoxon’s tests, respectively. B) NMDS of the Bray-Curtis dissimilarities (stress = 0.07, model fit = 99%) of AS and AB groups collected 12, 24, and 48 h after feeding. Ellipses represent confidence intervals of the distances calculated from each group’s centroids. ADONIS and ANOSIM tests support the contribution of each dependent variable (type of diet and time elapsed after feeding) in the distribution of microbial species. The different size of spheres is a function of the tridimensional perspective. C) Shared and unique microbial species between blood or sugar-fed adult mosquitoes. The intersections (connected dots) are color-coded by type of diet (Purples = Blood; Blues = Sugar), as are their correspondent bins in the histogram. The intersections sizes, representing the respective number of shared species between given groups are displayed in the histogram. Bins and dots of Species shared between different diets are black.

In addition to the rearing water used as an environmental control for the larval microbiome, the blood and sugar solutions used to feed the adults were also sequenced to assess the possibility that meals were the source of microorganisms colonization. In general, the number of reads attributed to microbial phylotypes was drastically lower in all control samples, corresponding to a decrease of 98.6% in assigned reads (268,671) compared to the mosquito samples (20,409,956 assigned reads). In the blood sample, more than 95% of reads were assigned to *Sus scrofa*, coinciding with the source of the blood used for the mosquitoes’ blood meals. A few viral reads were detected in the blood source and were assigned to the Taterapox virus, which was also present in a small portion of the microbiome of adult mosquitoes fed with blood (Figures 1B and S4), and indicative that females likely acquired these viruses during the blood meal. In the water-sugar solution used for feeding, the bacterial species *Microbacterium* sp. and *Pseudomonas fluorescens* were found with 666 and 441 reads, respectively. In the negative controls (blanks), *Methylobacterium* sp. was detected (34,651 reads; Supplementary Figure S4). This taxon was previously described as a contaminant in commercial kits commonly used for metagenomics DNA extraction (Salter et al., 2014). Together, the metagenomic analyses of experimental controls show that our analyses were not influenced by the microbial components previously present in the environment where larvae were reared, meal solutions, or in the DNA extraction kit.

### 2.3. Shifts in microbial composition are driven by hematophagy

The composition of the normalized data sets shows that Bacteria is the prevalent microbial domain in larvae and in the midgut of females of *A. aegypti* (approx. 99.6% Bacteria, 0.3% Fungi, 0.05% Viruses, 0.007% Archaea, Supplementary Figure S3), independently of the food source. Individually, the samples in the sugar-fed group displayed the most reads assigned to the Eukarya domain regardless of the time after feeding, unlike blood-fed adult mosquitoes and larvae (Supplementary Figure S3). Even though most of these reads were further confirmed to be remaining fractions of *A. aegypti’s* genome, 10% to 15% of these eukaryotic reads in mosquitoes of the AS group were composed of the fungal phylum Ascomycota (Figure 1A). Nevertheless, Proteobacteria was the phylum with the most significant portion of reads in all metagenomes analyzed, with relative abundances ranging from 45% to 95% of the microbiomes in individual samples (Figure 1A).

At the species taxonomic level, the top 50 microorganisms assigned to each sample are detailed in Figure 1B. A total of 802 microbial phylotypes were detected in the metagenomes with a per-group average of L4 = 301,5 ± 17.8; AB = 166.5 ± 93; AS = 18.5 ± 11. A shift in the microbiome composition was observed in the earliest hours after feeding the blood group (Figure 1). At 12h and 24h, the AB group displayed a dramatic increase in abundance of the phylum Proteobacteria, with the proliferation of Enterobacterales. At 48h post blood meal, the microbiome of the AB group exhibited yet another shift, and Enterobacterales was no longer detectable. However, at this time of the digestive period, a proliferation of the phylum Bacteroidetes was observed, with an average of 85% of the microbiome of female adults being composed of the flavobacterium *Elizabethkingia anophelis* (146,605 reads in the AB group at 48h of a total of 340,158 reads across all groups; Figure 1B). This shift is similar to what is observed in the AS group at 48h (44,348 reads assigned to *E. anophelis*), indicating a modulation of their proliferation capability triggered by the blood metabolism in the midgut environment (Figure 1). In the post-digestive period (48h), group AB presents a microbial composition similar to that observed in the AS groups at 12h and 24h, but with the presence of ascomycete fungi such as *Aspergillus flavus* and *Metarhizium majus* (with 12,508 and 10,260 reads attributed, respectively; Figure 1B).

### 2.4. Diversity of the microbiomes under different diets

In sugar-fed adults, the total number of microbial phylotypes observed (67 phylotypes) is relatively low throughout the digestive period. Blood-fed adults, conversely, display a high variation of observed phylotypes (average of 136 ± 18 phylotypes) during digestion, reaching the peak of 174 phylotypes 24h after the blood meal, as shown in Figure 2. The Simpson reciprocal index, however, indicates that the peak of microbial diversity in the sugar-fed group is reached at 12h, decaying in the next 36h (Figure 2A). The blood-fed group shows an inverse pattern, with its lowest microbial diversity at 12h after feeding and increasing diversity to higher levels, up to 48h (Figure 2A). The discrepancies between the indices indicate that a higher evenness (i.e., the lack of dominance of one or a few taxa) drives the increase of the diversity in sugar-fed adults when using indices that account for relative abundances of taxa, such as Simpson or Shannon. The contrary is also true for the blood-fed group, which has less evenness with a few highly abundant species (Figure 1A). Taking into consideration that the time for the digestion of blood in the midgut of *A. aegypti* mosquitoes lasts 30 to 40h (Downe, 1975; Felix et al., 1991), the overlapping of samples from blood and sugar-fed groups at 48h indicates that the microbial composition of both groups is similar after digestion. The major modulation of the microbiome is therefore triggered within 24h after feeding by the type of diet. Further compositional analyses in this work consider these digestive periods, and the 48h groups will henceforth be referred to as the “post-digestive” period.

Notably, sample ordinations show divergent beta-diversity patterns (Figure 2B). The microbiomes of sugar-fed mosquitoes at 12h and blood-fed at 12h and 24h are highly unique, forming separate clusters with distinct centroids (Figure 2B). Nevertheless, there is no clear distinction between the sugar-fed groups at 24h and 48h and the blood-fed group at 48h, which can be observed in the NMDS as the superposition of ellipsoids of these groups (Figure 2B, left panel). The scattered distribution of blood-fed individuals analyzed 48h after feeding caused its centroid to overlap with all individuals in the sugar-fed groups at 48h, as opposed to other groups of blood-fed individuals (Figure 2B), demonstrating a convergent shift back to similar compositions after 48h since their last meal, regardless of the diet. However, when individuals from groups AB and AS are analyzed independently of the time elapsed after feeding, the two groups are clearly separated (Figure 2B, right panel), thereby indicating that the type of diet is the best explanatory variable to the compositional microbiome differences (p < 0.001; ADONIS = 0.27), followed by hours post-feeding (p < 0.001; ADONIS = 0.16). These results corroborate previous studies demonstrating that meal sources may directly affect the mosquito microbial community (Almire et al., 2021; Gonzales et al., 2018; Muturi et al., 2018).

A high number of shared phylotypes (117 phylotypes) was observed in the microbiome of mosquitoes 12h and 24h after the blood meal (Figure 2C). This shared diversity of the midgut microbiome decreases significantly to 66 phylotypes after 48h of feeding in the AB group and even further when the AS group at 48h is considered (38 phylotypes; Figure 2C). The blood-fed group showed a lower number of unique microbial phylotypes independent of the time post-feeding, with 60 phylotypes at 12h, 105 phylotypes at 24h, and 16 at 48h. Such findings confirm the compositional narrowing of the microbial phylotypes after the blood meal. On the other hand, the sugar-fed group had a higher number of unique phylotypes (25 and 5 phylotypes, respectively) at 12h and 24h, as opposed to shared phylotypes (4 phylotypes shared between all AS groups; 3 phylotypes shared between AS individuals at 12h and 24h as well as between 12h and 48h) further confirming that the sugar feeding is associated with a more diverse microbial repertoire.

### 2.5. Microbial network interaction throughout digestion

Microbiomes are complex ecological communities that form interactive networks. To infer these interactions in the microbial community structure, we utilized graphs of microbial networks at the species level, aiming to detect taxa with significant co-occurrences (Figure 3). Sugar-fed mosquitoes present fewer microbial co-occurrences than blood-fed adults, but six clusters (S1 to S6) were significantly correlated (p < 0.05; R > 0.75; Figure 3A). In blood-fed mosquitoes, six clusters (B1 to B6) were also observed, and the densest network consists of three clusters, where B3 and B4 are composed exclusively of specific Enterobacterales phylotypes found in the AB group at 12h and 24h after feeding, respectively (Figure 3B). Intriguingly, a third cluster, B5, is composed of microbial taxa that are distantly related, unlike what was observed in the other clusters (Figure 3B). These results indicate that the structure of *A. aegypti’s* microbial community may not be driven only by the type of diet and digestion periods but also by taxon-specific interactions. The detection of several bacterial phylotypes belonging to the same genera, such as *Pseudomonas* spp. and *Elizabethkingia* spp. in clusters S4, S5, and B1, likely indicates that they are the same species colonizing the midgut of *A. aegypti* but methodological artifacts intrinsic to the sequence homology and similarity in closely related phylotypes may miss such nuances as intraspecific variation. Clusters marked with special characters (*, ◯, and #) are highly similar in composition between sugar- and blood-fed groups. These microbial phylotypes are detected only in the blood post-digestive period but are evenly distributed among individuals fed with sugar.

**Figure 3.**
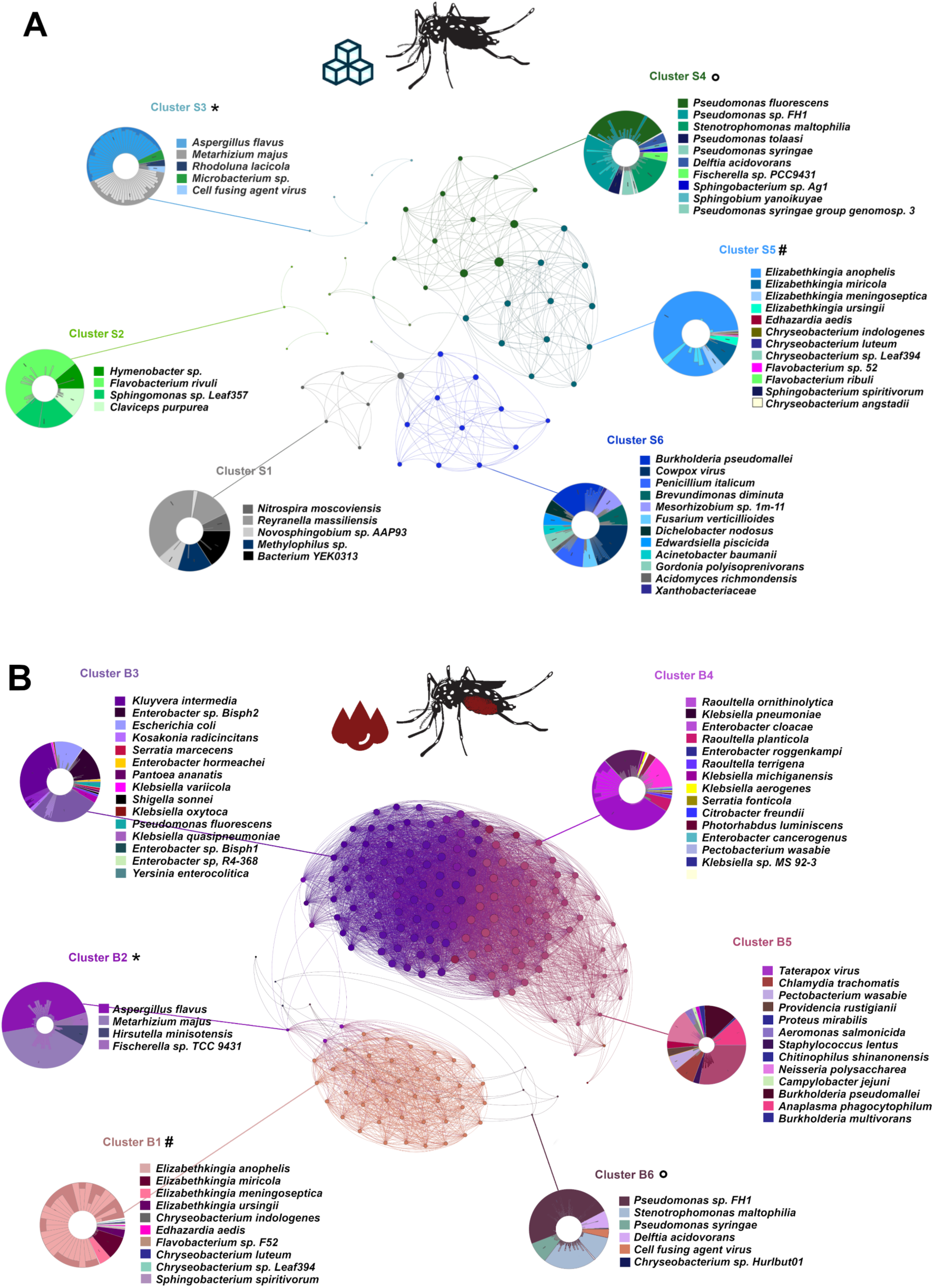
Co-occurrence networks representing the graph structure of the microbial community in adult mosquitoes fed with sugar (A) and blood (B), and their interactions. Graph vertices display microbial species, and edges represent their co-occurrences calculated with the Pearson correlation coefficient (r > 0.75; p < 0.05). Clusters marked with special characters (*,° and #) are similar in species composition.

Part of the microbial richness consistently observed in co-occurrence networks may be used as predictors of the diet and the time elapsed after a meal. This relationship can be observed in Figure 4A, which shows that *P. fluorecens*, *A. flavus*, and *M. majus* have the strongest association (p < 0.05) with the AS group at 12h. On the other hand, the microbiome of the AB group was dominated by the bacterial phylotype *Raoultella ornithinolytica* (169,307 reads) at 12h post-feeding, while *Kluyvera intermedia* (196,549 reads) is predominant at 24h post-feeding. The midgut of individuals metabolizing blood (12 and 24h) share a significant association with the phylotypes *Salmonella enterica*, *Enterobacter cloacae,* and other Enterobacterales phylotypes. After 48h, the relative abundance of *E. anophelis* is assigned to both groups of adult mosquitoes regardless of the diet, making this phylotype the most descriptive of the post-digestive state (Figure 4A). Further analysis of the abundance of Flavobacteriales and Enterobacterales revealed that both orders reach their peak richness opposite to all other taxonomic orders. The order Flavobacteriales showed less distinct peaks in the diversity distributions, whereas the Enterobacterales displayed only two well-defined richness peaks when no other microorganisms were detected. This finding indicates that the proliferation of these bacteria in the midgut of mosquitoes is mutually exclusive with the presence of other microbial components given the microenvironmental pressure presented by blood digestion (Figure 4B).

**Figure 4.**
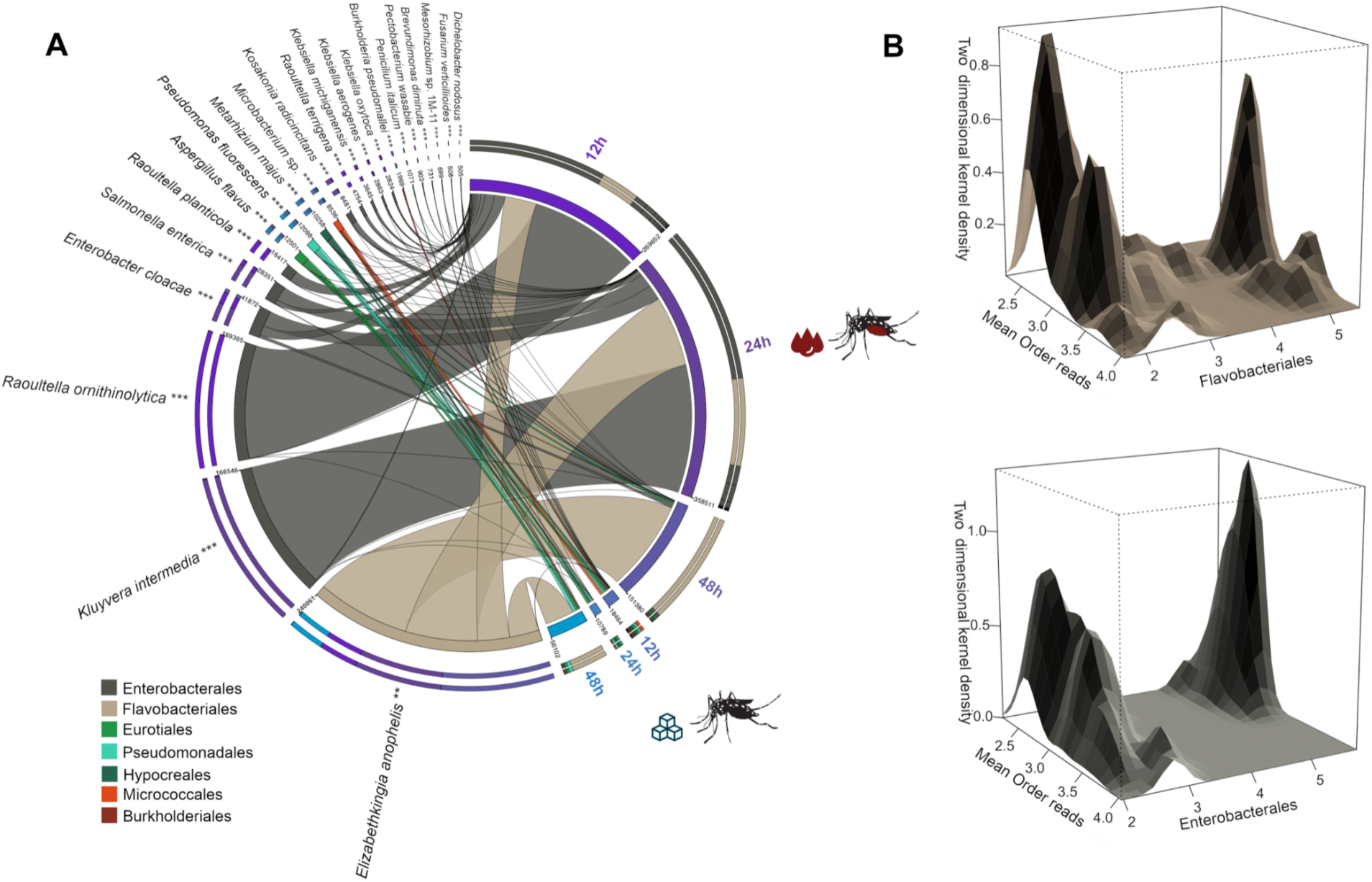
Microbial composition landscape in the midgut of adult mosquitoes. A) Occurrence of 22 microbial species predictive of digestive states (diet and digestion time). The width of the ribbons indicates the relative abundance in the linear scale. Ribbons link species to experimental groups (represented by the color of external circles; Blue = AS, Purple = AB) and are color-coded by the taxonomic order. B) Density estimates displaying the occurrence of reads identified in the taxonomic orders Flavobacteriales (top) or Enterobacteriales (bottom) in the x-axis, with the mean of reads attributed to other microbial taxonomic orders in the y-axis, transformed to the log10 scale.

The presence of *E. anophelis* in the post-digestion microbiomes of adult females is noteworthy, mainly because this gram-negative flavobacterium has been identified as the causative agent of multiple outbreaks in humans worldwide. The species has also been isolated from *Anopheles gambiae,* raising concerns about the possibility of its vectorial transmission (Chew et al., 2018; McTaggart et al., 2019; Perrin et al., 2017; Reed et al., 2020). Its pathogenesis is characterized by nosocomial bacteremia leading to sepsis in humans and has been associated with neonatal meningitis (Lau et al., 2016). Recent evidence suggests that this bacterium is also found in the saliva and salivary glands of *A. albopictus* (Onyango et al., 2021), indicating that transmission by mosquitoes is a possible route. Interestingly, in the same study, the colonization by *E. anophelis* was correlated with lower ZIKV titers in mosquito co-infection assays *in-vivo*. Although previous studies using metabarcoding have shown a high abundance of the flavobacterium *Chryseobacterium* sp. (Kim et al., 2005), our results do not indicate the prevalence of this phylotype in our datasets. However, *Chryseobacterium* and *Elizabethkingia* are closely related taxa, and the assignment differences found may be explained by the higher taxonomic resolution power of the WGS metagenomics, thus indicating that *Elizabethkingia*, in particular *E. anophelis,* is likely the dominant genus in the midgut microbiome of *A. aegypti*.

This evidence is further supported by the recovery of the metagenome-assembled genome (MAG) from the reads assigned to *Elizabethkingia* spp. The MAG is fragmented into 558 contigs with a total length of 4,667,330 bp (N50 = 32,698 bp). A ribosomal multilocus sequence typing (rMLST) resulted in 38 loci (out of 53) with 92% support for the MAG assignment to *E. anophelis*. Further fIDBAC analysis provided the identification of the MAG as *E. anophelis* based on >98% average nucleotide identity (ANI). Furthermore, the comparative phylogenomic analysis of 1,254 single-copy orthologs of 185 genomes corroborated the grouping of the recovered MAG into *E. anophelis* species clade (Supplementary Figure 5). This study reports the first evidence of the colonization of *A. aegypti* by *E. anophelis*. Our results show that both sugar and blood diets interfere with the proliferation of this bacterium in the midgut. The relationship between *E. anophelis*, blood meal ingestion, antiviral activity, and pathogen transmission is worth further investigation.

### 2.5. Functional profiling of the microbiomes throughout digestion

The functional profile of metagenomes was characterized using the SEED database (Overbeek et al. 2014) to analyze microbial composition’s impact on the midgut’s functional diversity. Results indicate that, similarly to what was observed in the microbial compositional analyses, the functional profiles of midguts of blood-fed mosquitoes are strikingly different at 12h and 24h compared to other adult experimental groups (Figure 5). Most of the metabolic classifications are pathways related to the cell wall and ultrastructure (4,395 reads), nucleotide metabolism (44,694 reads), and stress response (1,861 reads in the AB groups). However, 12 hours after feeding, fewer genes are related to motility and chemotaxis than in other groups (30,194 reads). After 48 hours, the functional profile of the AB group shifts back to a state that resembles those observed in sugar-fed groups (Figure 5), further showing that the shifts in microbial composition and function are concomitant.

**Figure 5.**
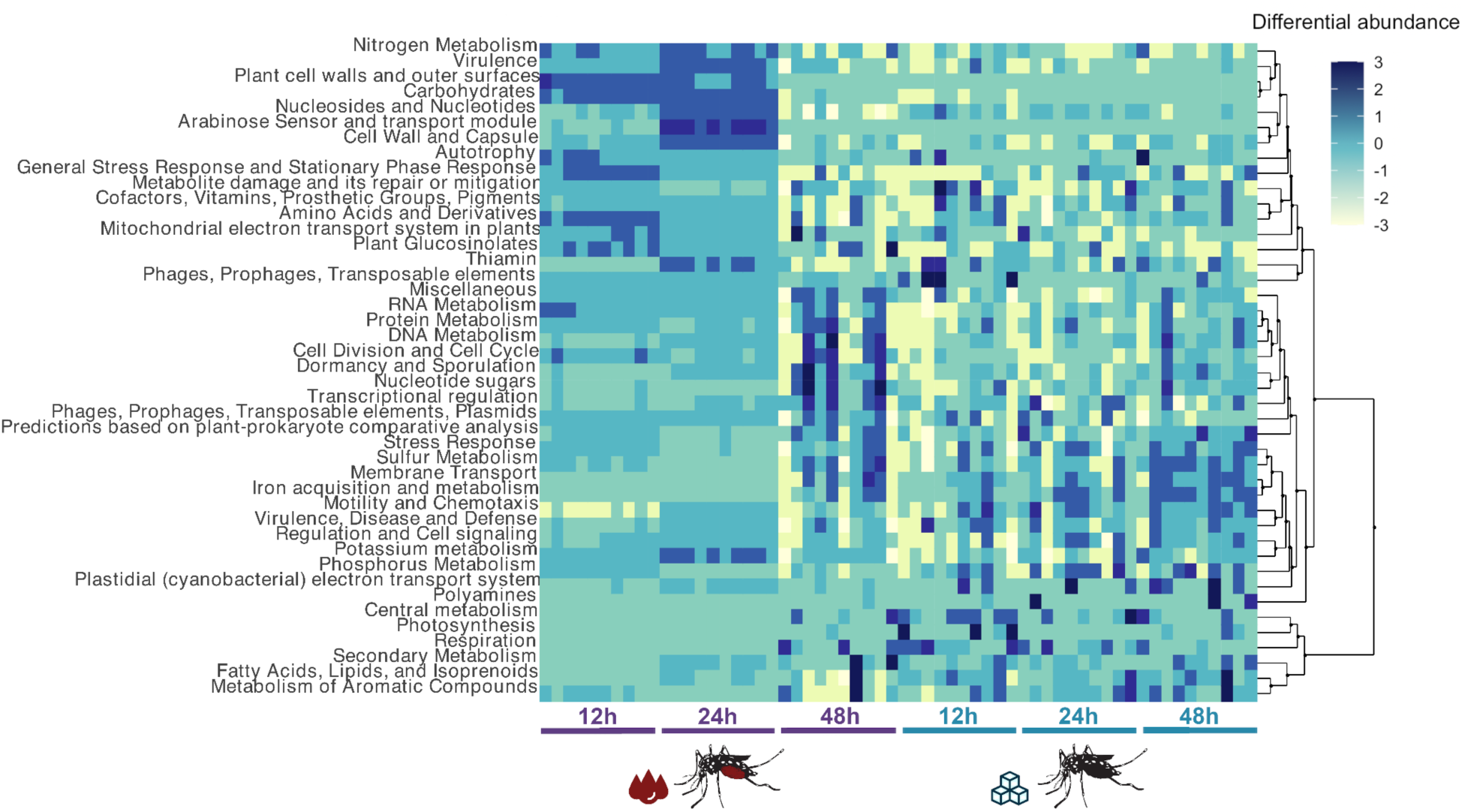
Functional classification of microbial reads in the midgut of adult females fed with blood and sugar. The heatmap was generated using z-score transformed values acquired in the metabolic pathway enrichment using SEED pathways functional classes in the metagenomes analyzed. Functional pathways are organized by hierarchical clustering.

A principal coordinate analysis (PCoA) demonstrated that reads assigned to carbohydrate metabolism (209,958 reads), amino acids and derivatives (127,903 reads), and virulence (100,710 reads) are responsible for the uniqueness observed in the functional profile of blood-fed mosquitoes at 12 and 24h (Figure 6A). Such singularity displays a strong positive correlation with the proliferation of Enterobacterales (Figure 6B), namely *S. enterica* (28,351 reads; Rho = 0.83), *Ko. radicincitans* (8,480 reads; Rho = 0.73), *Kle. aerogenes* (2,863 reads; Rho = 0.74) and *Klu. intermedia* (196,549 reads; Rho = 0.70). These bacterial phylotypes are the most significantly correlated with the functional uniqueness in the AB group at 12 and 24h, as shown in Figure 6C. Additionally, the principal coordinate that best explains the variability of all SEED metabolic pathways (PC1 = 58,2%) has a significant negative correlation with the relative abundance of Enterobacterales (Figure 6D), thereby indicating that these bacteria are likely responsible for narrowing the metabolic spectrum in the mosquito midgut during the digestion of blood, and are in agreement with the functional diversity analysis shown in Figure 5. This trend can be better observed in Figure 6A (upper panel), which shows a higher number of metabolic pathways in the midgut of mosquitoes in groups AB at 12 and 24h than in other groups (Figure 6E, top). Yet, the functional variability (Evar; Figure 6E, bottom) observed in group AB at 12 and 24h is lower when compared to all other groups, showcasing that, even though a large number of pathways is detected, most of the reads are concentrated in the three pathways aforementioned. These results indicate that the blood meal triggered the proliferation of specialized opportunistic Enterobacterales, also suggesting that these bacteria play a more significant role than previously thought in the digestion of blood in the mosquito’s midgut. This hypothesis corroborates previous work describing the main association of the mosquito microbiome with gene expression related to metabolic and nutritional pathways (Hyde et al., 2020). Different Enterobacteria species also have been associated with an increased digestive capability of fructose in *A. albopictus* (Scolari et al., 2019), but this is the first study that describes their potential for partaking in blood digestion in hematophagous insects. It is possible to speculate that these Enterobacteria may be related to all digestive processes of the mosquito *in vivo*, albeit not associated with non-natural processes, such as the digestion of sucrose utilized in our analyses.

**Figure 6.**
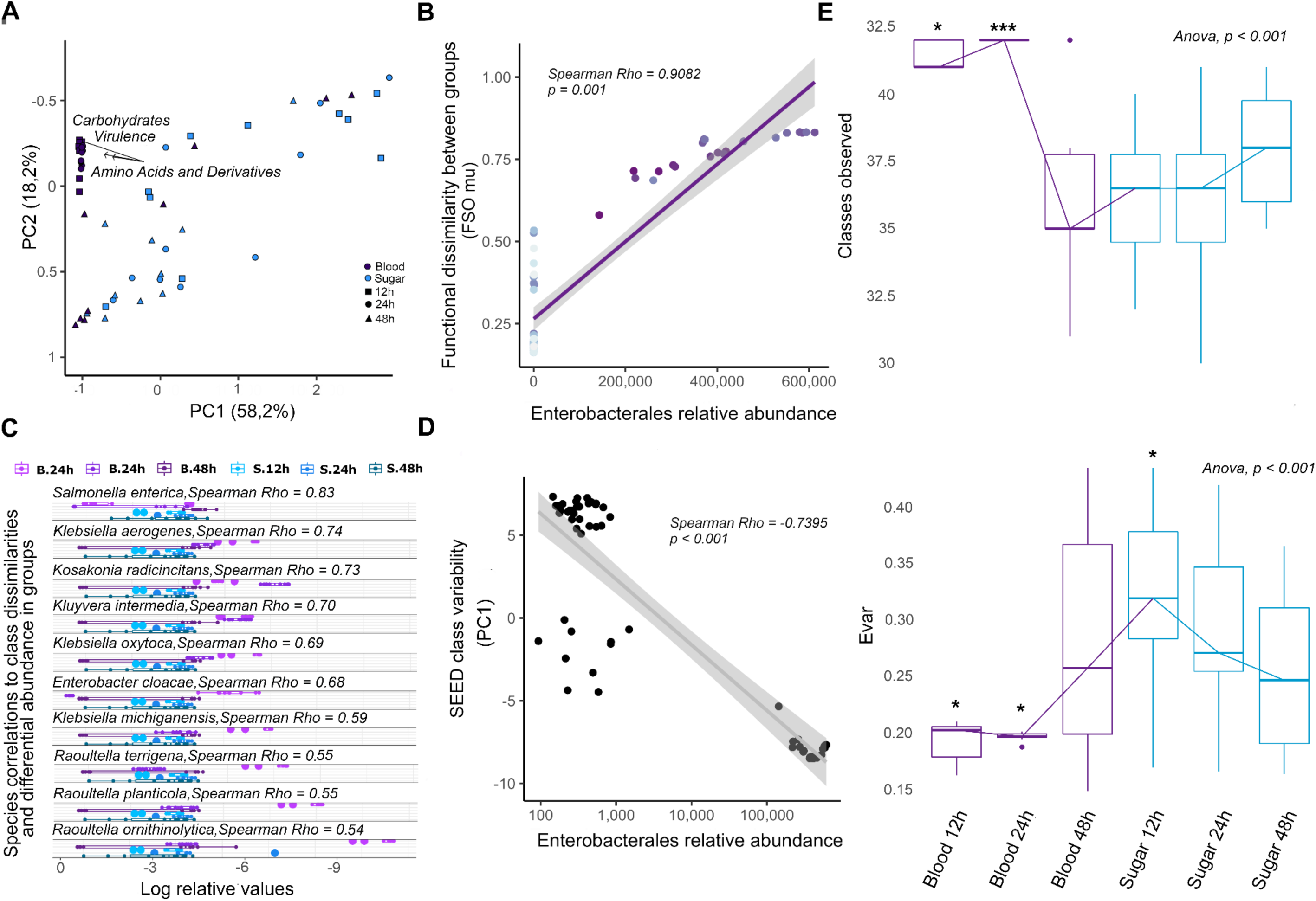
Functional diversity of the midgut microbiome in adult mosquitoes. A) PCoA of Bray-Curtis dissimilarities (PC1 = 58.2% vs PC2 = 18.2%) displaying SEED pathways responsible for the sample ordination. Colors represent different diets and shapes represent hours post-feeding. B) Fuzzy set ordination using a generalized linear model (GLM) to display the correlation between a functional dissimilarity matrix and the relative abundance of Enterobacteriales. C) Differential occurrence of Enterobacteriales species classified as predictors for the blood diet in log-relative scale and their individual correlations with the functional dissimilarities. D) Scatter plot using GLM to display the correlation between the principal explanatory coordinate (PC1) and the relative abundance of Enterobacterales. E) Boxplots showing the distribution of observed SEED pathways and their variability indices (Evar). Medians are indicated by the trend line, and the global and pairwise significances are assessed with ANOVA and Wilcoxon’s tests, respectively.

To further analyze the modulation of significantly correlated Enterobacterales phylotypes (*S. enterica*, *Ko. radicincitans*, *Kle. aerogenes,* and *Klu. intermedia*) in the digestive period, the relative abundance of reads assigned to each phylotype and principal functional pathways (Rho > 0.70; carbohydrate metabolism, amino acids and derivatives, and virulence) are shown for the AB group at 12 and 24h in Figure 7. It is possible to observe an overlap of metabolic pathways and microbial abundance of specific Enterobacterales, reiterating the strong relationship between the taxonomic and functional classification in the midgut of blood-digesting mosquitoes. Considering the virulence pathway, it is possible to detect the presence of functional classes related to multiple antimicrobial resistances, notoriously multi-resistance efflux pumps (12,324 reads), and fluoroquinolone resistance (4,743 reads; Figure 7C). The detection of commensal microorganisms that present antimicrobial resistances has already been reported in *A. aegypti’s* midgut with culture-dependent methods (Hyde et al., 2019), but this is the first metagenomic evidence of the presence of a putative resistome associated with the microbiome in these mosquitoes. Under the carbohydrate metabolism pathway, most of the SEED classes are related to the utilization and anabolism of sugars, including maltose and maltodextrin (9,527 reads), and serine utilization in the glyoxylate cycle (15,409 reads). The enrichment of these genes may indicate an increase in the metabolic demand caused by the proliferation of Enterobacteria, in particular that of *Klu. intermedia* after blood meals in *A. aegypti*. Nonetheless, previous studies demonstrate that the consumption of dextrose may increase adult mosquitoes’ lifespan (Alvarado et al., 2021; Carter and Evans, 2005; de Campos et al., 2016; Posidonio et al., 2021; Singh et al., 2004), suggesting that the blood meal may indirectly influence the hosts’ fitness by driving the proliferation of beta-hemolytic Enterobacteria which may be reducing or increasing the availability of macro-nutrients to the host, but further studies are needed to confirm this hypothesis. Lastly, the amino acids and derivatives pathway include functional classes related to glycine and serine metabolism (10,368 reads), as well as biosynthesis (16,643 reads) and degradation (12,018 reads) of methionine (Figure 7C). These pathways have key enzymes for the digestion of blood, which is the case of the glycine and serine utilization pathway, detected in the microbiome of adult mosquitoes. The silencing of one of the pivotal enzymes in the latter pathway (serine transferase, SMHT) caused the formation of clots of non-digested blood in female mosquitoes’ midguts and a phenotype of ovarian underdevelopment (Li et al., 2019), corroborating with our results and strengthening the hypothesis of a nutritional symbiosis between Enterobacteria and *A. aegypti*.

**Figure 7.**
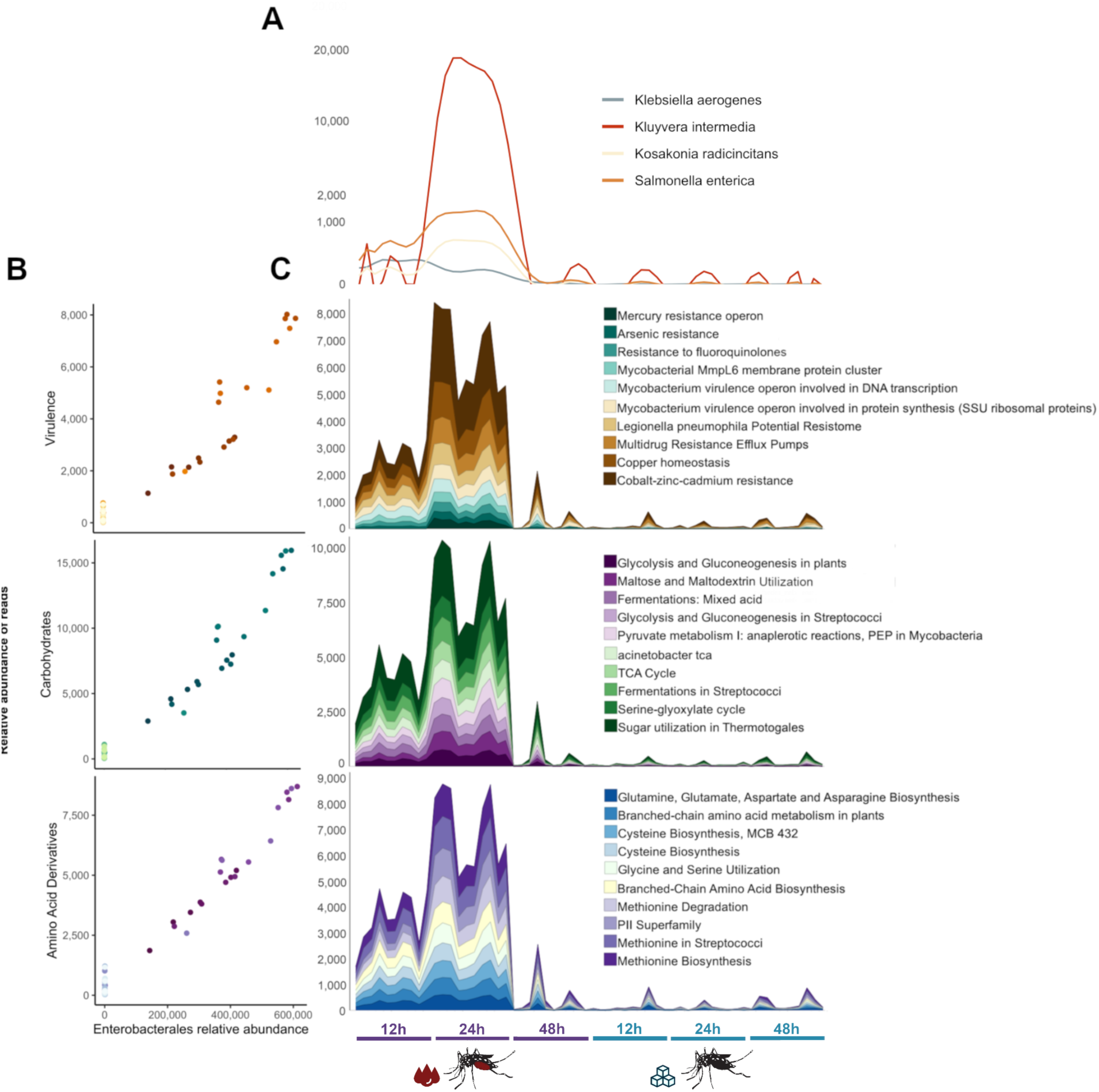
Distribution of the relative abundance of reads attributed to highly explanatory SEED pathways, and Enterobacteria in the metagenomic datasets. A) Polynomial distribution of reads attributed to Enterobacteria species correlated to the variability of functional profiles. These distributions are displayed in quadratic scale in the y axis. B) Scatter plots displaying the comparison between the distribution of reads attributed to Enterobacteria *sensu latu* and to the Amino Acids and Derivatives, Carbohydrates Metabolism and Virulence pathways. The stacked line charts display the distribution of the respective SEED pathways and its uncollapsed branches in quadratic scale.

## 3. Concluding remarks

Associations between microorganisms and insect hosts have contributed to a better understanding of physiology, metabolism, and their relation to vector capacity and transmission of insect-borne diseases. In this work, we sequenced individual microbiomes of larval and female adults of *A. aegypti* fed with blood and sugar, demonstrating that transstadial transmission of microorganisms from larval to adult stages is not prominent in *A. aegypti*. We also observed that, after digestion (48h), no significant difference was observed in the microbiome of blood-fed or sugar-fed individuals. In fact, significant and transient changes in microbial composition, diversity, and putative function are mainly driven by blood meal and last for the digestion period in adults. Furthermore, we observed a significant association of Enterobacterales phylotypes, especially of *Klu. intermedia* with blood digestion in the midgut. Hematophagy is a habit intimately associated with the mosquito’s reproductive cycle, and the Enterobacteria proliferation in response to the blood meal stimulus is likely associated with important metabolic and physiological changes required to cope with oxidative stress and blood digestion. The functional characterization inferred from the metagenome showed that potential changes in pathways follow the microbial compositional shifts that occur after a blood meal intake. The digestive process is capable of modulating the presence of fungi and prominent bacteria, such as *E. anophelis*, which are consistently observed colonizing the midguts of individual mosquitoes during the post-digestive period and may act as symbionts of *A. aegypti*. Both, *Klu. Intermedia* and *E. anophelis* are promising candidates for further assessment of their physiological impacts on the mosquito host and could potentially serve as control agents for both the vector population and the transmission of arboviruses.

## 4. Material and methods

### 4.1. Mosquito rearing and feeding

*A. aegypti* Singapore strain was reared in a Climatic Test and Plant Growth chamber (Panosnoic MLR-352H) at a temperature of 28 ± 1°C, relative humidity of 80 ± 5%, and a photoperiodic regime of 12:12 h (light:dark). Mosquito breeding followed a previously established protocol (Zhang H et al., 2023). In brief, 2–4-week-old *Ae. aegypti* eggs were hatched in sterile water using a vacuum for 15 minutes. Newly hatched L1 larvae were bred in a plastic bowl with a density of 2.5 mL/larva in sterile water. Mosquito larvae were fed with a mixture of fish food (TetraMin Tropical Flakers)/brewer’s yeast (yeast instant dry blue/Bruggeman) at a ratio of 2:1. The feeding regimen was as follows: 25 mg (day 1), 32 mg (day2), 56 mg (day 3), 130 mg (day 4), 200 mg (day 5), and 100 mg (day 6). Pupae were collected and placed into a 17.5 x 17.5 x 17.5 cm cage (BugDorm-4S 1515) supplied with one vial of sugar and one vial of sterile water, both of which were replaced twice per week.

### 4.2. Experimental design

For sugar-feeding (AS group) mosquitoes, 80 newly emerged *Ae. aegypti* mosquitoes (40 females and 40 males) were maintained in a 17.5 x 17.5 x 17.5 cm cage (BugDorm-4S 1515) supplied with one vial of sugar and one vial of sterile water. For blood-fed (AB group) mosquitoes, 80 newly emerged *Ae. aegypti* mosquitoes (40 females and 40 males) were maintained in a 17.5 x 17.5 x 17.5 cm cage (BugDorm-4S 1515) supplied with one vial of sugar and one vial of sterile water for three days to allow maturation before being fed with *Sus scrofa domesticus* blood using an artificial membrane feeding system (Hemotek Ltd, UK, or Orinnotech, Singapore). For each group, three subgroups of ten adults were separated based on time elapsed after their last blood (AB) or sugar (AS) meal: 12h, 24h, and 48h. Additionally, ten individuals in the L4 larval stage were analyzed to compare the two developmental stages and assess whether the larval microbiome is transmitted to adults. One sample of the larvae rearing water (W), one sample of the distilled water source used to prepare the water-sugar solution (DW), one sample of the water-sugar solution itself (WS), and one sample of the *Sus scrofa domesticus* blood (B) were analyzed to assess their contribution to the microbiome of mosquitoes. Additionally, three non-mosquito (NM) samples for DNA extraction reagent control were also sequenced, corresponding to the three extraction protocols we used: (NM-1) one negative control for the kit DNeasy Blood and Tissue (Qiagen) used for mosquito DNA extraction, (NM-2) one negative control for the kit DNeasy Blood and Tissue (Qiagen) used for DNA extraction of minipig blood and (NM-3) one negative control for the kit DNeasy PowerWater (Qiagen) used for the DNA extraction of water samples. Thus, a total of 77 samples (Supplementary Table 1) were sequenced, processed, and analyzed with the same workflow.

### 4.3. DNA extraction and sequencing

Midguts of adult female mosquitoes were dissected in ice-cold PBS using a stereoscope (Olympus SZ61) and kept in 300 µL of Phosphate Buffered Saline (PBS) 1X, pH 7.4 (Gibco – ThermoFisher). The midguts and larvae were individually macerated with an electric tissue grinder (VWR International), and the homogenates were used for DNA extraction following the insect tissues protocol of the DNeasy Blood and Tissue (Qiagen) kit, according to the manufacturer’s instructions. The mini pig blood sample was extracted with the same kit but following the specific protocol for blood DNA extraction. We followed the standard protocol of the kit DNeasy PowerWater (Qiagen) for the water sample extractions. Negative control DNA extractions followed the respective protocols, but no sample was added. DNA yield quantification was performed with the Qubit^™^ 1X dsDNA HS Assay Kit (ThermoFisher) in a Qubit 2 fluorometer (ThermoFisher), and DNA integrity was assessed on a Bioanalyzer 2100 system (Agilent) using the High Sensitivity DNA Kit (Agilent). The total DNA for each sample was fragmented using the ultrasonicator Covaris S220 (Covaris Inc.), and the fragments were separated by size in a Pippin Prep electrophoretic system (Sage Science) with 2% agarose gel. Fragments of 300 to 450 bp were collected and purified with Agencourt AMPure XP magnetic beads (Beckman Coulter). Libraries were then built with the Accel-NGS 2S Plus DNA Library Kit (Swift Biosciences), following the manufacturer’s protocol. All libraries were indexed with the 2S Dual Indexing Kit (Swift Biosciences), quantified with Quant-iT^TM^ Picogreen^®^ (Invitrogen), and validated by qPCR with the KAPA SYBR^®^ FAST qPCR kit (Kapa Biosystems). Equimolar quantities of each indexed library were pooled for multiplex sequencing on the HiSeq 2500 (Illumina Inc.) platform, with a 251 bp paired-end protocol. Sequencing was performed at the Singapore Centre for Environmental Life Sciences Engineering, Nanyang Technological University (Singapore).

### 4.4. Processing of sequenced datasets

The raw fastq sequencing files were trimmed for both adapter and low-quality sequences using cutadapt v. 1.15 (Martin, 2011). A maximum error rate of 0.2 was allowed to recognize and remove adapters. A quality cutoff of Q20 was used to trim low-quality ends from reads before adapter removal. High-quality reads were aligned against the complete *A. aegypti* genome (GCA_002204515.1) to filter out the host reads. Mapping was carried out using *Bowtie2* (Langmead and Salzberg, 2012) with selective parameters for a high sensitivity rate, and reads were filtered with SAMtools (Li et al., 2009). The remaining fraction of reads was subsequently translated in six frames and aligned against the NCBI NR protein database with RapSearch2 v. 2.15 (Zhao et al., 2012) using default parameters. The number of reads generated and analyzed is listed in Supplementary Table S2.

### 4.5. Taxonomic and functional assignment

After importing alignment results into MEGAN 6 v. 6.18.8 (Huson et al., 2016), we performed taxa assignment with strict parameters of the Lowest Common Ancestor (LCA) algorithm, considering the read length generated for each sample with the following settings: Max Expected = 0.01, Top Percentage = 10.0, Min Support = 25, Min Complexity = 0.33, Paired Reads = On. Next, we individually normalized all metagenomes to the dataset with the smallest number of reads of the 70 experimental or seven control samples to obtain the representative relative abundances of assigned microbial taxa. The functional profiles for the different microbiomes were assessed by assigning enriched genes identified to functional classes using the SEED hierarchy (Mitra et al., 2011) database. Results were visualized in a heatmap adjusted to a z-score scale, with hierarchical grouping of classes.

All reads assigned to *E. anophelis* in the metagenomic analysis were extracted with an in-house script and used as input in SPAdes v. 3.15.2 to assemble the genome (Prjibelski et al., 2020). The resulting MAG was evaluated using QUAST v. 5.0.0 (Mikheenko et al., 2018) and ribosomal multilocus sequence typing (rMLST) was performed using the speciesID tool available in the public databases for molecular typing and microbial genome diversity PubMLST (Jolley et al., 2012; release 2023-03-10) to confirm the metagenomic assignment. The average nucleotide identification (ANI) was performed with fIDBAC (Liang et al., 2021). A pangenome approach was also conducted to identify orthologs for tree reconstruction. Briefly, protein datasets of 183 annotated, non-redundant genomes of *E. anophelis* available on NCBI (Supplementary Table S2 for accession numbers; downloaded on 12/21/2021) were used as an input for the software Orthofinder v. 2.5.4 (Emms and Kelly, 2019). The genome of *Elizabethkingia miricola* (GCF_001483145.1) was used as an outgroup. A total of 1,254 single-copy orthologs present in all samples were assigned by Orthofinder and individually aligned with MAFFT v. 7 (Katoh and Standley, 2013). The concatenated alignment of orthologs was then used as input to IQ-TREE v.1.6.12 (Nyugen et al., 2015) to perform the best-fit model search (JTT+F+R4) with ModelFinder (Kalyaanamoorthy et al., 2017) and reconstruct the maximum likelihood tree with 10,000 replicates of ultrafast bootstraps (Minh et al., 2013).

### 4.6. Diversity estimations and statistical analyses

Species diversity analyses using the Simpson Reciprocal (Simpson, 1949) and Shannon-Weaver Indexes (Shannon, 1948) were performed in MEGAN 6, while the package vegan v. 2.5-6 (Oksanen et al., 2019) was used to generate the chao1 richness index (CHAO et al., 1990) using “species” level for Bacterial, Fungal and Viral taxa of NCBI taxonomy. Analysis of variance (ANOVA) was performed for each method with 1000 permutations, and pairwise group significance was assessed with Wilcox’s post hoc test. To display diversity as a function of the sampling size, rarefaction curves were computed with functions of the package iNEXT v. 2.0.20 (Hsieh et al., 2016), with parameters of interpolation and extrapolation for Hill numbers (Hill, 1973). Calculation of non-metrical multidimensional scaling (NMDS) was also performed in vegan, using Bray-Curtis dissimilarity (Bray and Curtis, 1957) and calculating ellipsoids with confidence intervals based on the centroids for each experimental group. For the statistical interpretation of the results, multivariate analyses with distance matrices (ADONIS) and analysis of similarities (ANOSIM) were employed with 1000 permutations. The distribution of functional classes was calculated with the Bray-Curtis dissimilarity using a principal coordinates analysis (PCoA). To further investigate the functional diversity, a fuzzy set ordination (FSO) was employed using the functional dissimilarities, using the package fso (Roberts, 2008). Visualizations were plotted using the package ggplot2 v. 3.3.0 (Wickham, 2016).

### 4.7. Distribution of significant species

Linear regressions were directly employed in the dispersion of quantitative variables to assess the correlation of relative species abundance to the observed functional variability in the metagenomes. To test whether microbial species originate from the same time-diet distribution, we used a phyloseq (McMurdie and Holmes, 2013) implementation of the Kruskal-Wallis non-parametric test coupled with decision trees, in a model adapted from Torondel et al. (2016). Briefly, the p-values were calculated and corrected for multiple testing using familywise error rate for each pair “phylotype -experimental group”. The significance was based on the corrected threshold of p < 0.05. Significant species most frequently predicted were then assigned importance in the Random Forest classifier based on the mean decrease in accuracy of their classification (Breiman, 2001). To map the probabilistic density of abundant bacterial orders, their kernel bivariate densities (Venables and Ripley, 2002) were estimated as a function of the median distribution of reads assigned to other microbial orders, and their relationships are visualized in logarithmic scale. Additionally, co-occurrence networks were estimated for the microbiome of adult mosquitoes to detect potential microbial interactions related to different diets and the time of the digestive process. The networks were generated based on an adjacency matrix with a significance threshold p <= 0.05 for each microbial association with a bivariate Pearson correlation coefficient > 0.75 (Pearson, 1895). The attribution of edges was automated with the module iGraph (Csardi et al., 2006), removing vertices with null edges, thereby facilitating the visualization with the software Gephi (Bastian et al., 2009). As the input to these analyses, we generated a new abundance matrix based on the presence of species in at least 10% of the samples, with a minimum of 200 reads, removing likely contaminations and rare taxa to avoid algorithm miscomputations. All packages were accessed with *in-house* scripts written in R v.3.6.3 and are available upon request (R Core Team, 2013).

## Data availability statement

Sequencing data and metagenomes of *A. aegypti* will be available at Short Read Archive under the BioProject accession number PRJNA1065965. The specific BioSample accession numbers for each mosquito are found in Supplementary Table S3.

## Acknowledgements

JFMS is a doctoral fellow in the program ‘International Max Planck Research School: Principles of Microbial Life’, supported by funding from the Max Planck Institute for Terrestrial Microbiology, Marburg, Germany. This study was partially funded by Fundação de Amparo à Pesquisa do Estado do Rio de Janeiro to ACMJ (grant E26/211.473/2021).

## Author contributions

A.C.M.J., Y.C. and S.C.S. designed the study; Y.C., F.G.G and X.H. performed the breeding and feeding experiments as well as the midgut dissection. E.L.O. processed the samples and performed DNA extractions. D.I.D.-M. conducted the library preparation and sequencing. B.N.V.P., A.C.M.J. and S.C.S processed the raw sequencing data. J.F.M.S., B.N.V.P. and A.C.M.J. analyzed the metagenomic data. J.F.M.S. performed the statistical analyses. J.F.M.S and A.C.M.J wrote the manuscript with the input from B.N.V.P., D.I.D.-M., Y.C., and S.C.S.

## Competing financial interests

The authors declare no competing financial interests.

The authors declare that the manuscript has not been published previously and is not under consideration for publication elsewhere. All authors approved the submission of the article in the present form.

