## Supplementary Figure for "The Interplay Between Nutrition and the Dynamics of the Midgut Microbiome of the Mosquito *Aedes aegypti* Reveals Putative Symbionts"

**Supplementary material**

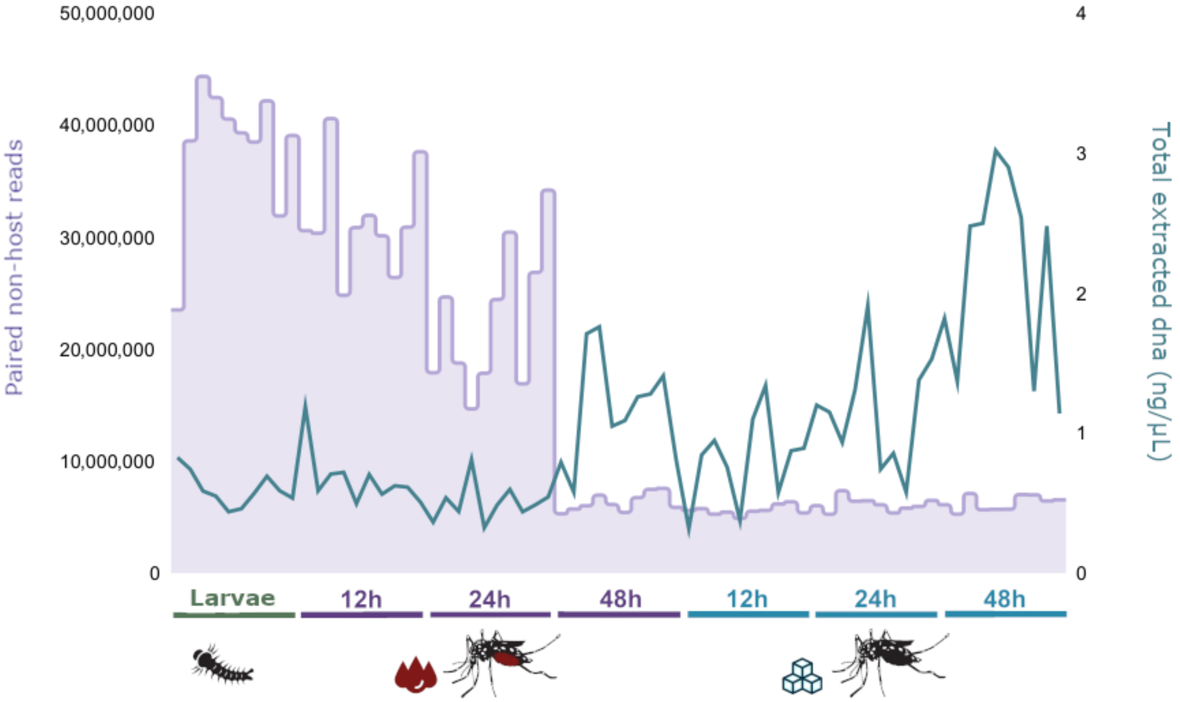


**Supplementary figure S1.** Total number of non-host reads used for the microbial taxonomic assignment (y-axis, left) and total DNA yield (y-axis, right) per individual sample (x-axis) of each experimental group.

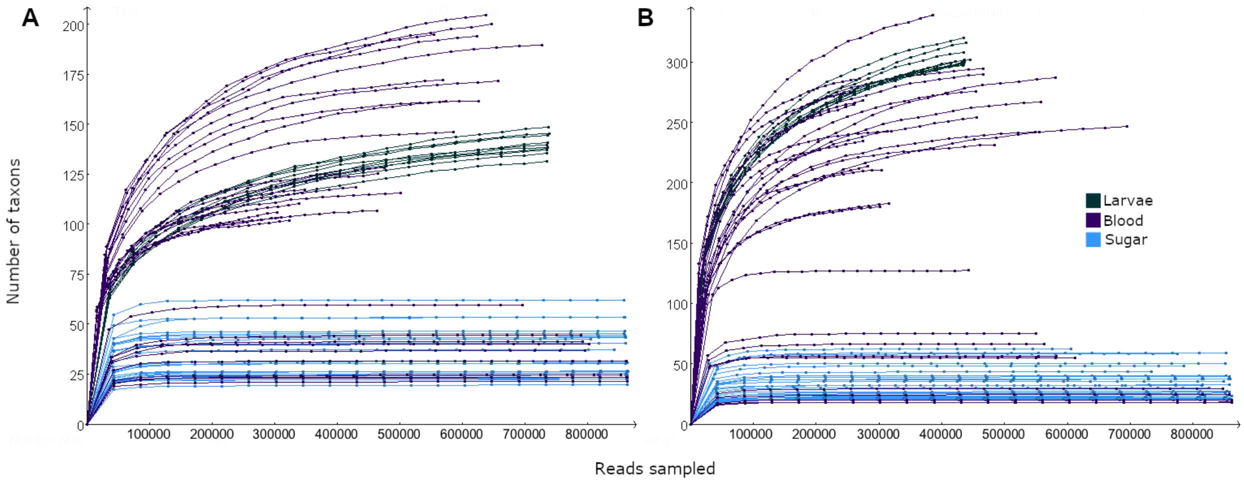


**Supplementary figure S2.** Rarefaction curves of the normalized number of reads as a function of the richness of identified microbial genera (A) and species (B). The taxonomic assignment was based on the NCBI *taxonomy* database. Curves are color-coded by the experimental groups of samples.

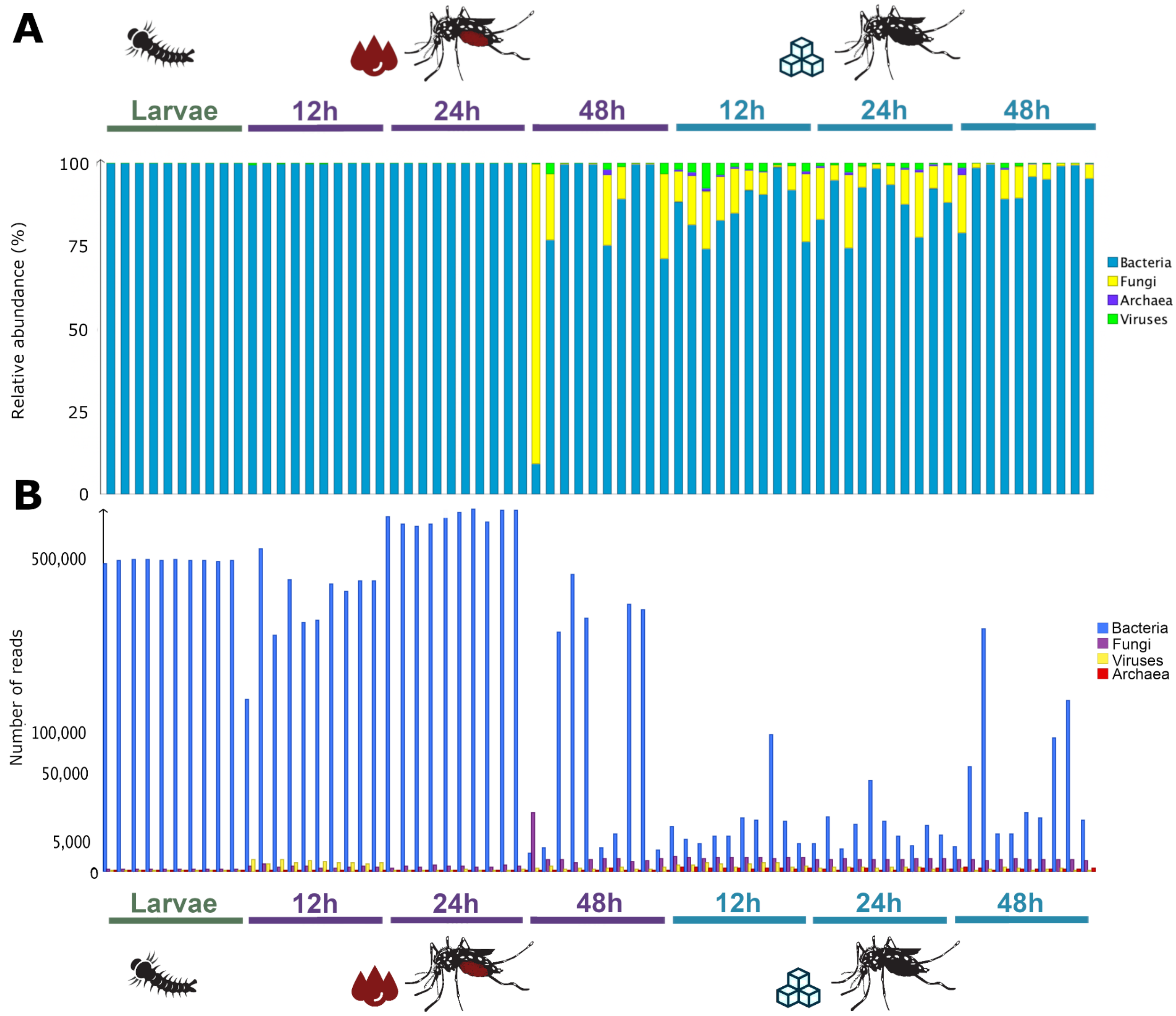


**Supplementary figure S3.** Microbiome composition in Superkingdom taxonomic level for Bacteria, Virus, Archaea and the Kingdom Fungi (classification according to NCBI), showing the normalized number of reads attributed to each taxon in the y axis in percentual (A) and quadratic (B) scales.

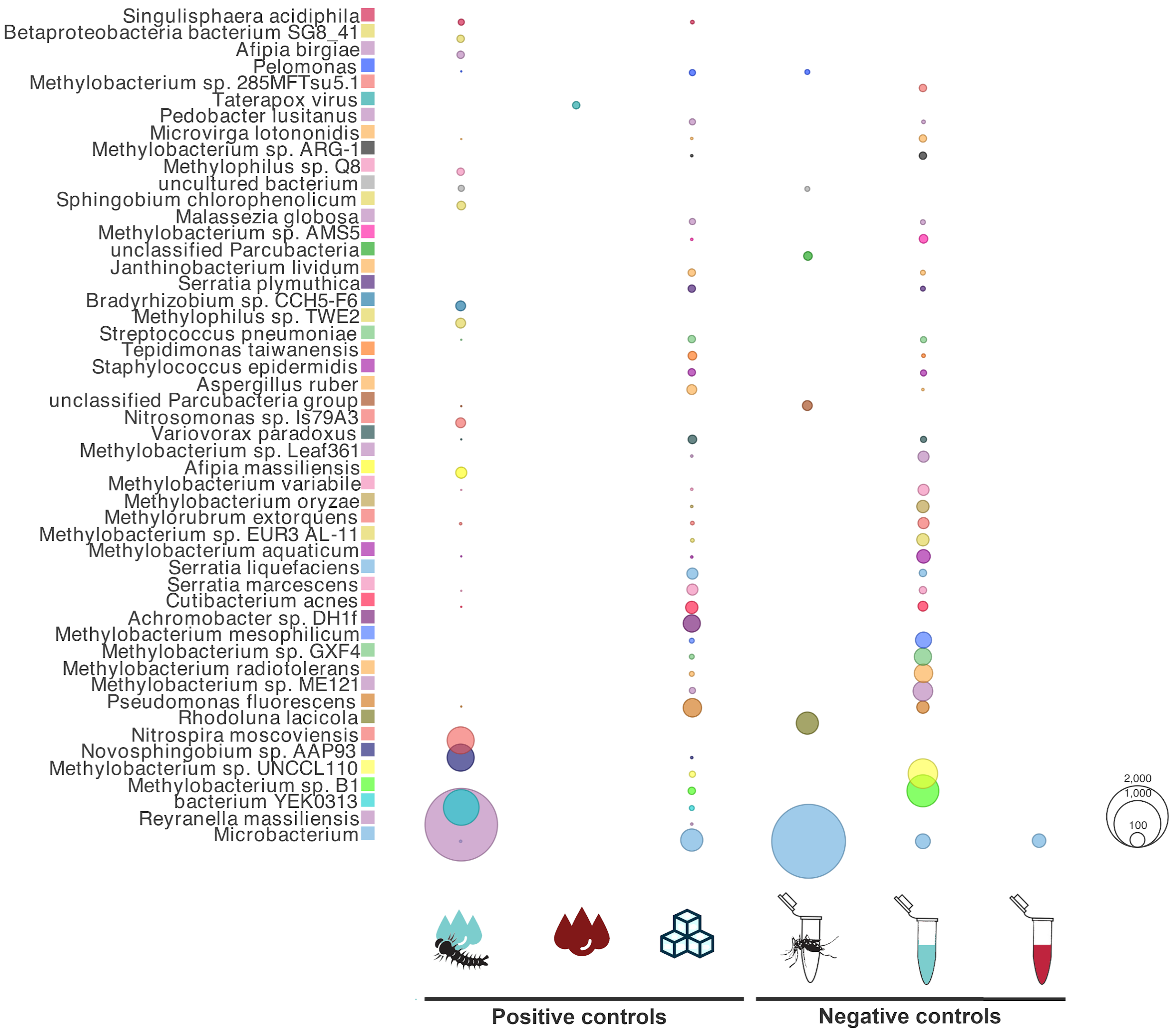


**Supplementary figure S4.** Microbial species detected in the positive or negative controls, corresponding, respectively to the experimental controls (samples CLW, CB and CW) and to the DNA extraction kit reagent blanks (samples NCM NCB and NCW). Bubbles represent the relative abundance of reads in quadratic scale.

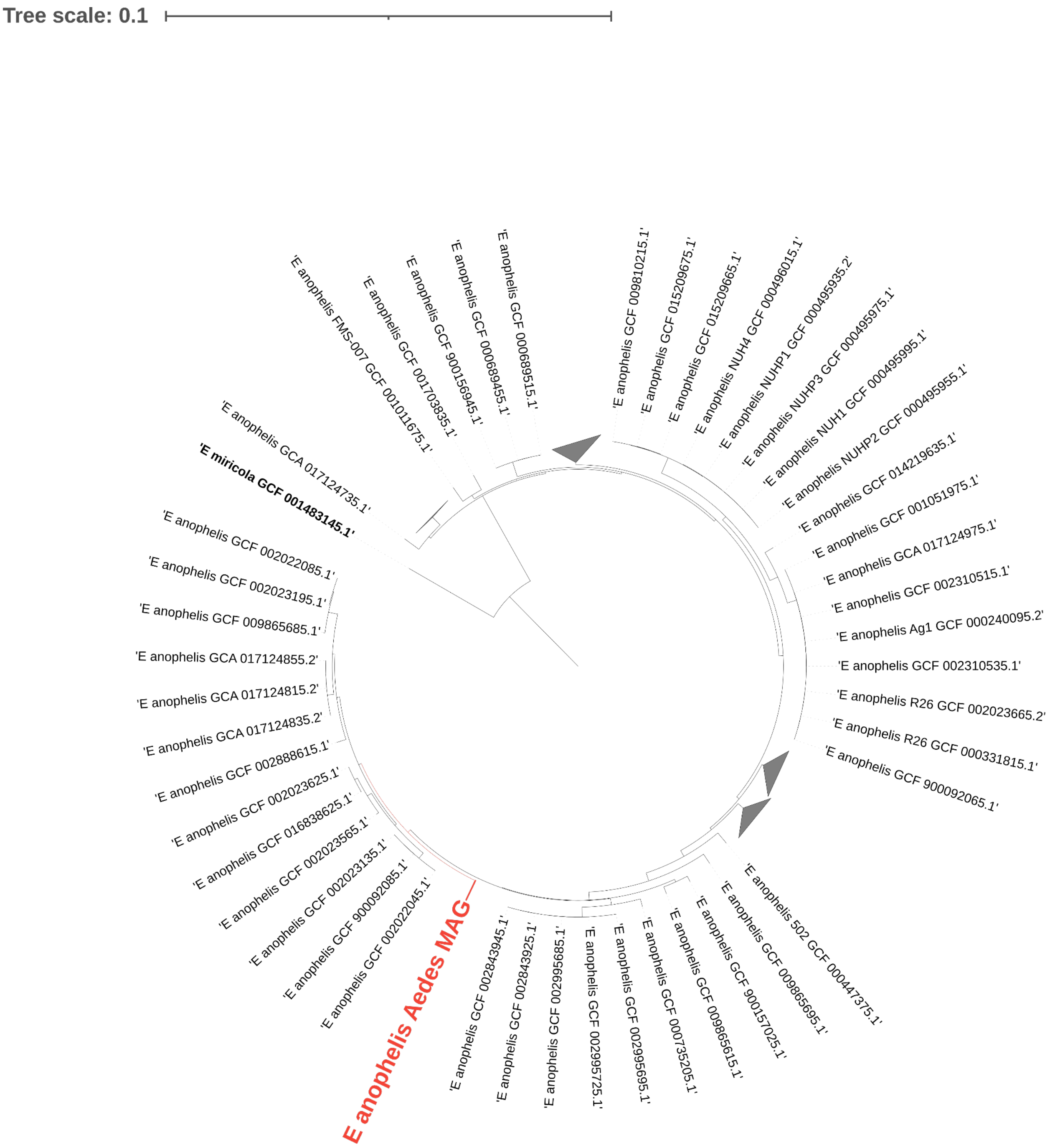


**Supplementary figure S5.** Maximum likelihood tree of the concatenated alignment of 1254 single copy orthologs from 185 of *Elizabethkingia*. spp. showing the putative positioning of the *Elizabethkingia anophelis* genome obtained from *Aedes aegypti*, displayed in red. The tree was rooted using *E. miricola* and non-informative *E. anophelis* clades were collapsed for visualization.
